# Inhibitory effect of an anti-prokineticin-1 antibody on liver metastases in mice injected with human colorectal cancer cell lines

**DOI:** 10.1101/2021.08.05.455319

**Authors:** Hiroko Kono, Takanori Goi, Hidetaka Kurebayashi, Katsuji Sawai, Mitsuhiro Morikawa, Kenji Koneri

## Abstract

Controlling hematogenous metastases is an effective treatment strategy for colorectal cancer. Multidisciplinary treatment for colorectal cancer has made great strides, and molecularly-targeted drugs have greatly improved the prognosis of patients. However, currently accepted molecularly-targeted therapeutic agents require concomitant use with anticancer agents. Thus, new molecularly-targeted drugs need to be developed. The prokineticin family of angiogenic factors has the potential of becoming target molecules. Among them, prokineticin-1 (PROK1) is involved in the promotion of angiogenesis, tumor growth, and liver metastases in colorectal cancer. We manufactured our own anti-PROK1 antibody and verified its effect in inhibiting liver metastases and prolonging survival. The method involved creating liver metastasis model mice using human colorectal cancer cell lines. These mice were divided into anti-PROK1 antibody administration and control groups. Mice were treated intraperitoneally with antibodies or phosphate-buffered saline (control) every 3 days. The number of liver metastatic lesions and survival time of each group were compared. The number of metastatic lesions decreased, and survival time was significantly prolonged in the antibody-treated group. Furthermore, using microarray and immunostaining in both groups, we confirmed the effect of administering the anti-PROK1 antibody on the oxidation, reduction, and apoptotic processes, and cell division of tumors, and that alterations were suppressed in 72.1% of the genes examined. The expression of transforming growth factor-β (TGF-β), a tumor suppressor gene, was increased. The increased expression of TGF-β via PROK1 antibody administration may suppress the cancer cell proliferation ability, leading to liver metastasis suppression and prolonging the survival time of mice.

## Introduction

Colorectal cancer is a highly prevalent malignancy worldwide, including in Japan [1-4]. The prognosis of colorectal cancer at an early stage is favorable, but the prognosis of unresectable, advanced colorectal cancer is not yet satisfactory. According to statistics of the Ministry of Health, Labor, and Welfare in Japan, the number of deaths from colorectal cancer continues to increase, exceeding 50,000 in 2016.

While there are various metastasis modes in colorectal cancer, such as lymph node, peritoneal, and hematogenous metastases, most patients face poor prognosis due to hepatic and other hematogenous metastases [5-7]. Recently, the prognosis of patients with colorectal cancer has improved due to great progress in multidisciplinary treatment. However, there is currently no treatment method that is capable of significantly improving the prognosis of patients when distant metastasis or recurrence is observed.

Angiogenic growth factors are important, and several such factors are thought to be involved in the development of colorectal cancer [8-10]. The prokineticin family of proteins, which we have focused on and studied throughout the years, is one family of angiogenic factors. It consists of two types of proteins: prokineticin-1 (PROK1) and PROK2. PROK1 is expressed in normal endocrine tissues of the adrenal gland, ovaries, and testes, and promotes the growth of vascular endothelial cells under hypoxic conditions. However, PROK1 is not homologous to vascular endothelial growth factor (VEGF) and is completely different from the known VEGF family of proteins [11].

To date, our laboratory has reported the involvement of PROK1 in tumor growth, angiogenesis, and infiltration in colorectal cancer through in vitro and in vivo experimental systems [12-14]. In this study, we used the anti- PROK1 antibody that was prepared in our laboratory and demonstrated its metastasis inhibitory effect on human colorectal cell lines using a liver metastasis mouse model.

## Materials and Methods

### Confirmation of PROK1 expression in human colorectal cancer cell lines

Human colorectal cancer cell lines (HCT116, HT29, DLD-1) were cultured at 37 °C under 5% CO_2_ for 3 days using RPMI medium with 10% fetal bovine serum. They were then frozen with an optimal cutting temperature compound, dissected into 4-μm sections with a microtome, and stained overnight at 4 °C using an anti-PROK1 antibody (Novus Biologicals, Littleton, CO, USA).

### Liver metastasis mouse model

Human colorectal cancer cells (1 × 106) were injected under the spleen capsule of male SHO mice to prepare the liver metastasis mouse model. Antibody administration and control groups were prepared for each colorectal cancer line (n = 5). In the antibody treatment group, the anti- PROK1 antibody (500 μg) was administered intraperitoneally the day before tumor cell injection under the spleen capsule and every 3 days after injection. In the control group, phosphate-buffered saline was administered intraperitoneally in the same manner. The anti-PROK1 antibody was originally prepared by our department as described previously [15].

The number of metastatic regions in the liver and the survival times of each group were compared. Mice were examined daily for their general condition and signs of moribund behavior. The mice were considered moribund when they could no longer reach out to water and/or food and were euthanized within 4 h of reaching moribund status. Survival curves were established using the Kaplan-Meier method, and a significant difference was determined at p < 0.05 using the log-rank test.

### Microarray

Using ISOSPIN Cell & Tissue RNA (Nippon Gene, Yoyama, Japan), total RNA was extracted from the primary lesions and metastatic regions in the liver and spleen of mice, and a microarray was performed. In the microarray, 24,351 genes were analyzed using the Clariom S Array, and human GeneChip (Thermo Fisher Scientific, Waltham, MA, USA).

### Immunohistochemical staining of liver metastases

Liver metastatic tissue was sliced at 10 μm thickness with a microtome and stained overnight at 4 °C with an anti-Ki67 antibody (Novus Biologicals). The mean numbers of positive cells at 400⍰ magnification were compared between the two groups, and the Mann-Whitney U test was used to determine any significant difference, which was set at p < 0.05. All statistical analyses were performed using EZR software [16].

## Results

### Immunostaining of human colorectal cancer cell lines

Cytoplasm was stained with the anti-PROK1 antibody in the HCT116, HT29, and DLD-1 cell lines (Fig 1). This demonstrated the presence of PROK1 in these lines.

**Fig 1.**
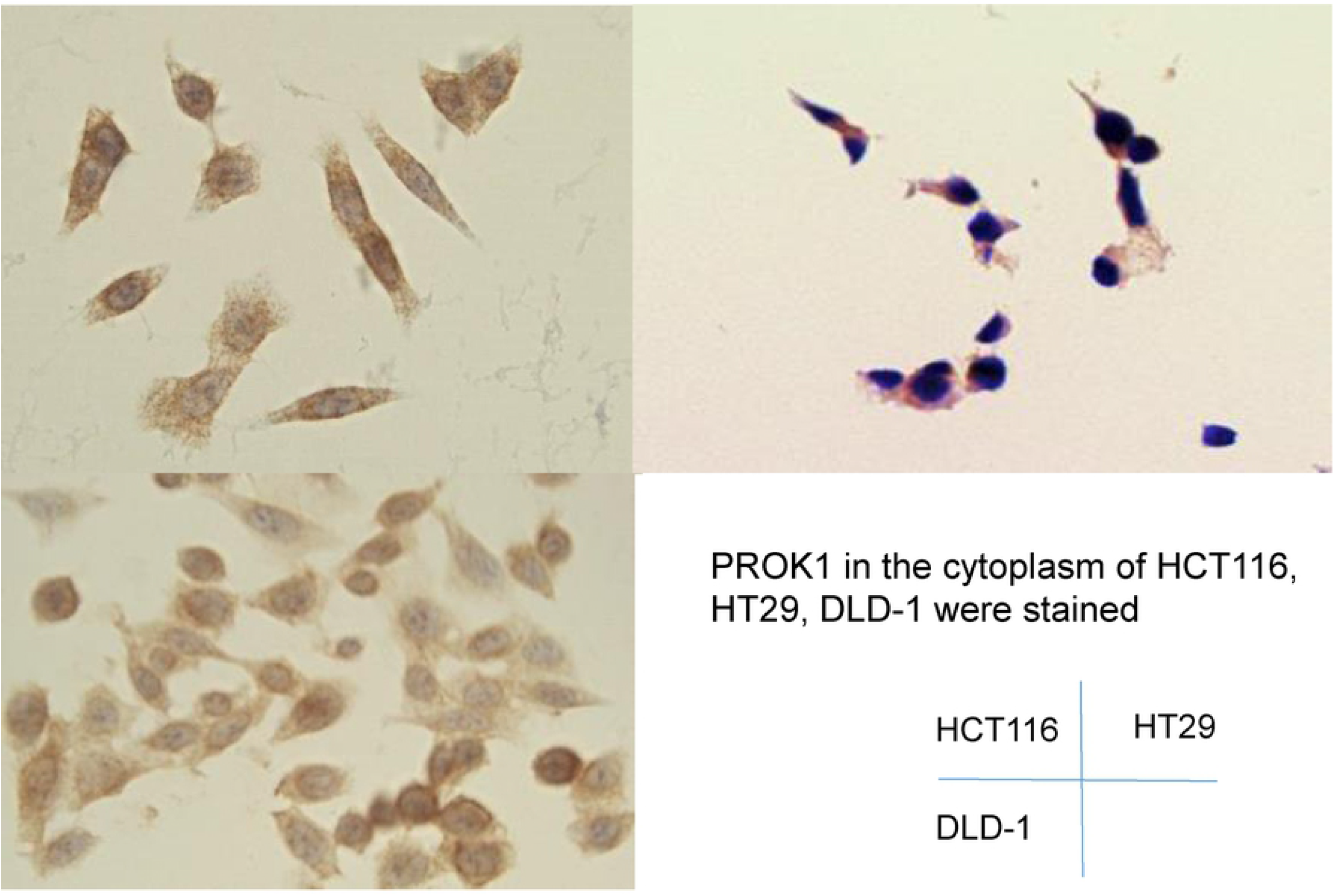
Immunohistochemical staining of colorectal cancer cell lines. Prokineticin-1 (PROK1) is stained in the cytoplasm of HCT116, HT29, and DLD-1 cells.

### Reduction of liver metastasis

Two weeks after injection of each colorectal cancer cell line, we confirmed that liver metastases were formed in all mice (Fig 2A). We removed the livers and confirmed the number of metastatic regions in each liver (Fig 2B). The median numbers of these regions were: HCT116 (control group: 95; antibody group: 68); HT29 (control group: 70; antibody group: 60); and DLD-1 (control group: 9; antibody group: 2). These results indicated that there were fewer liver metastases in the antibody-administered than in the control group.

**Fig 2.**
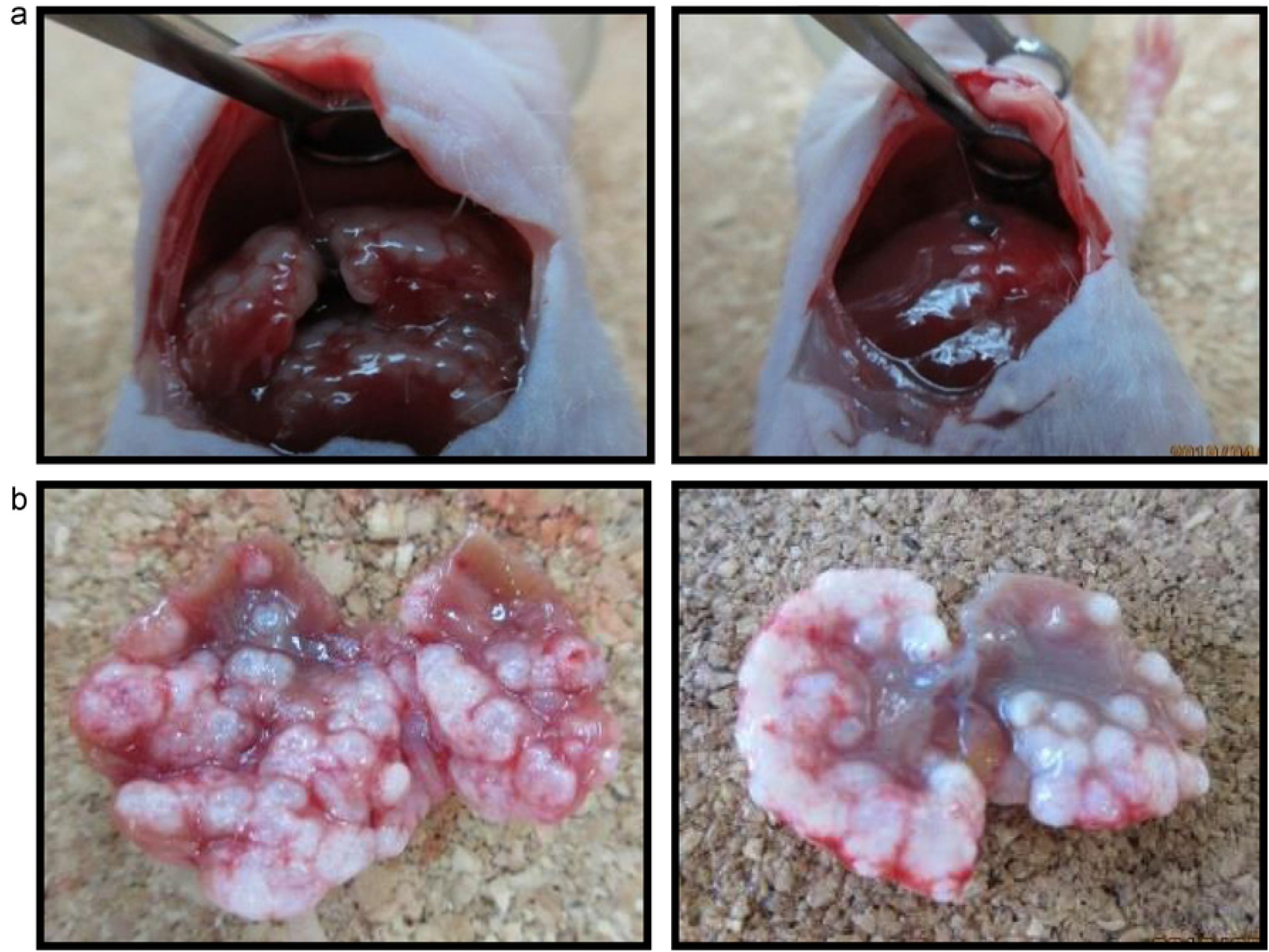
Images of liver metastases. A. Liver metastasis 2 weeks after spleen injection of the HCT116 cell line. Left: control group, Right: antibody group. B. Liver removed at the time of death. Left: control group, Right: antibody group.

### Extension of survival time

We established survival curves for each group and compared the median survival times: HCT116 (control group: 33 days; antibody group: 26 days; p = 0.00885); HT29 (control group: 28 days; antibody group: 46 days; p = 0.0279); and DLD-1 (control group: 28 days; antibody group: 13 days; p = 0.00249). The p-value for the survival time of mice (n = 15) was p = 0.0273. The survival time was significantly prolonged in the antibody-administered group (Fig 3).

**Fig 3.**
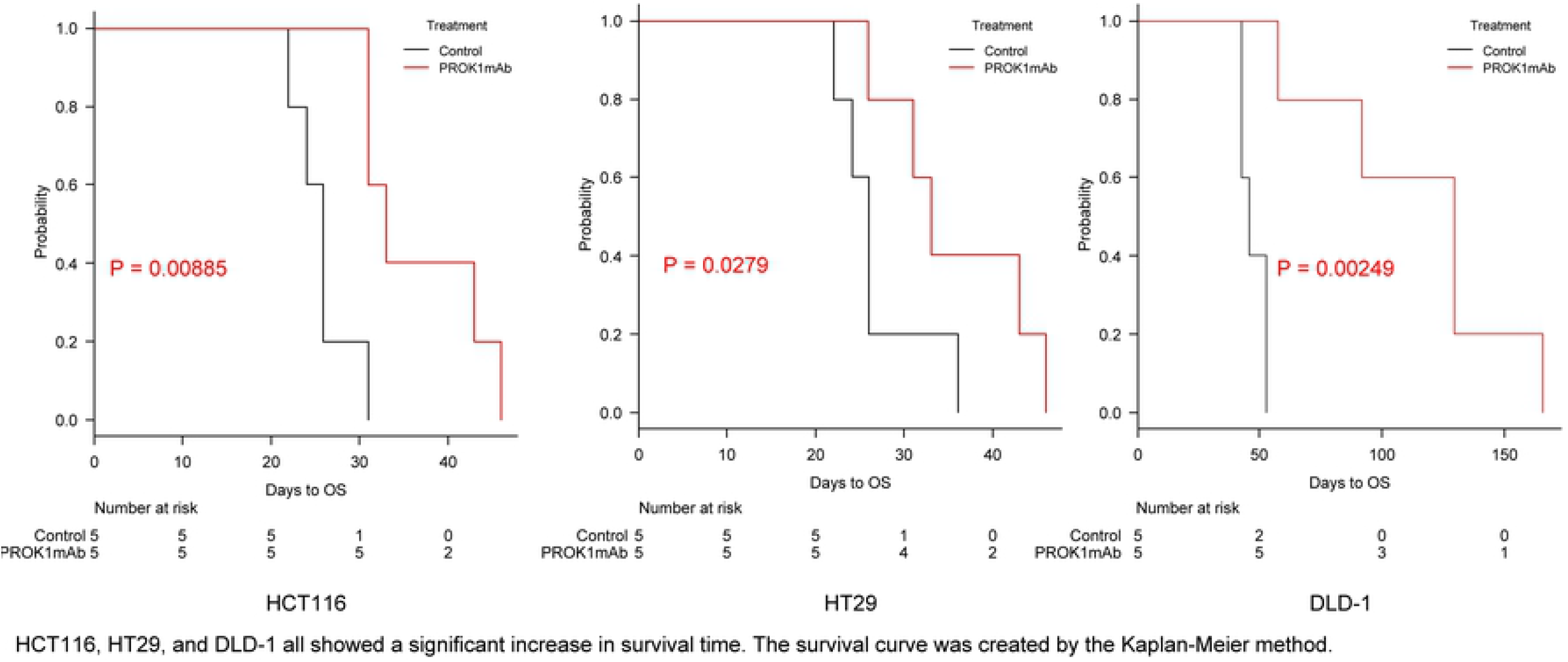
Comparison of survival times. Mice injected with HCT116, HT29, and DLD-1 cells all show a significant increase in the survival time when treated with the anti-prokineticin-1 antibody. The survival curve was created by the Kaplan-Meier method.

### Microarray

The heat map showed that genetic changes were inhibited in the antibody-treated mice (Fig 4). Specifically, the microarray revealed that changes were inhibited in 72.1% of the 24,351 analyzed genes. In addition, when the functions of genes that were significantly changed upon antibody administration (genes that had increased or decreased expression more than two-fold due to antibody administration) were analyzed using DAVID functional annotation, we found that the expression of genes involved in oxidation-reduction and apoptotic processes, and cell division, were significantly changed (p < 0.05) (Fig 5A). Among the altered genes, some were involved in the p53 cascade. In addition, there was an increase in the expression of transforming growth factor-β (TGF-β), which is a tumor suppressor gene (Fig 5B).

**Fig 4.**
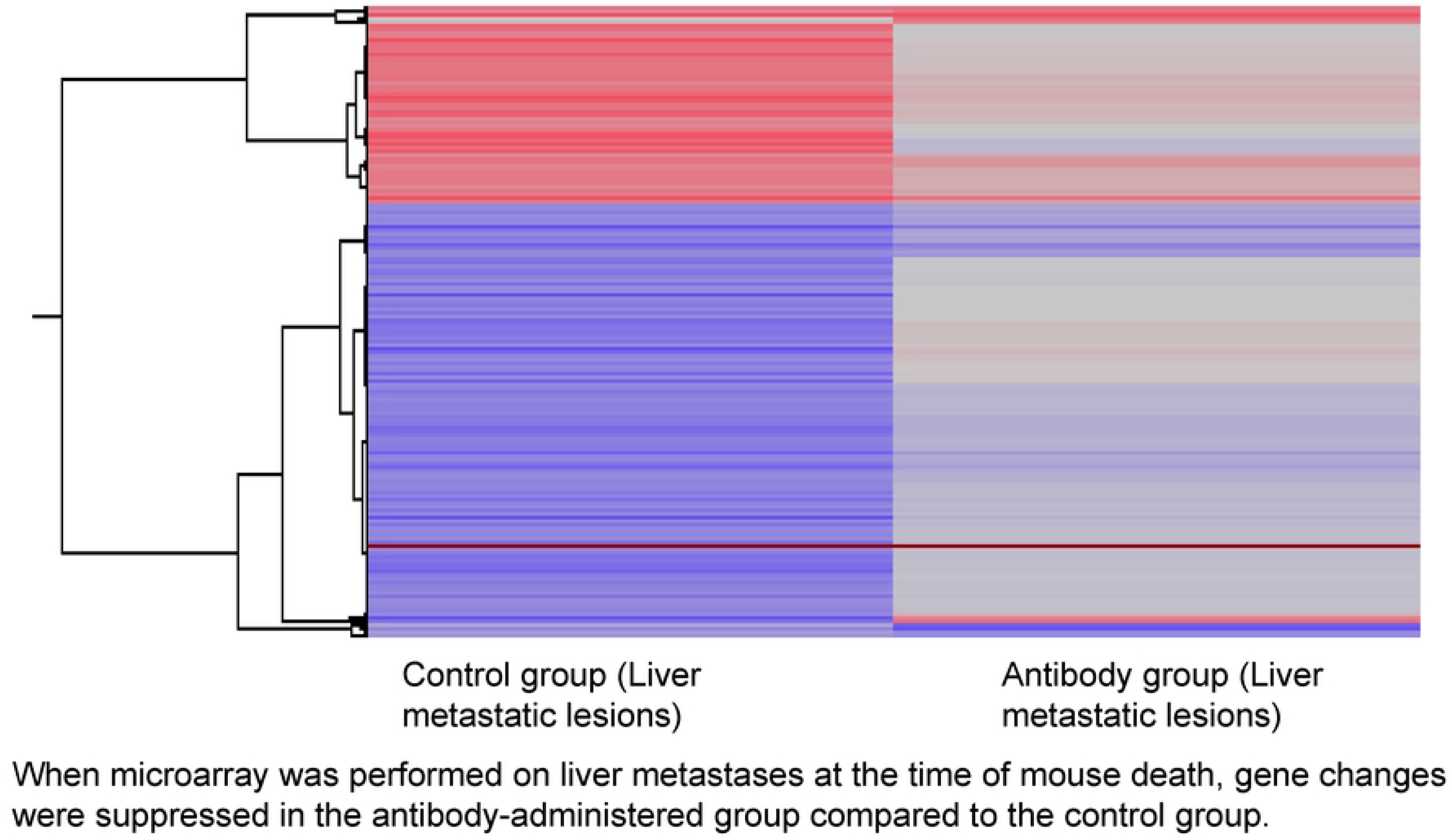
Microarray heatmap. A microarray was performed to detect liver metastases-related genetic changes at the time of death. This map shows that genetic changes are suppressed in the anti-prokineticin-1 antibody-administered group compared to the control group.

**Fig 5.**
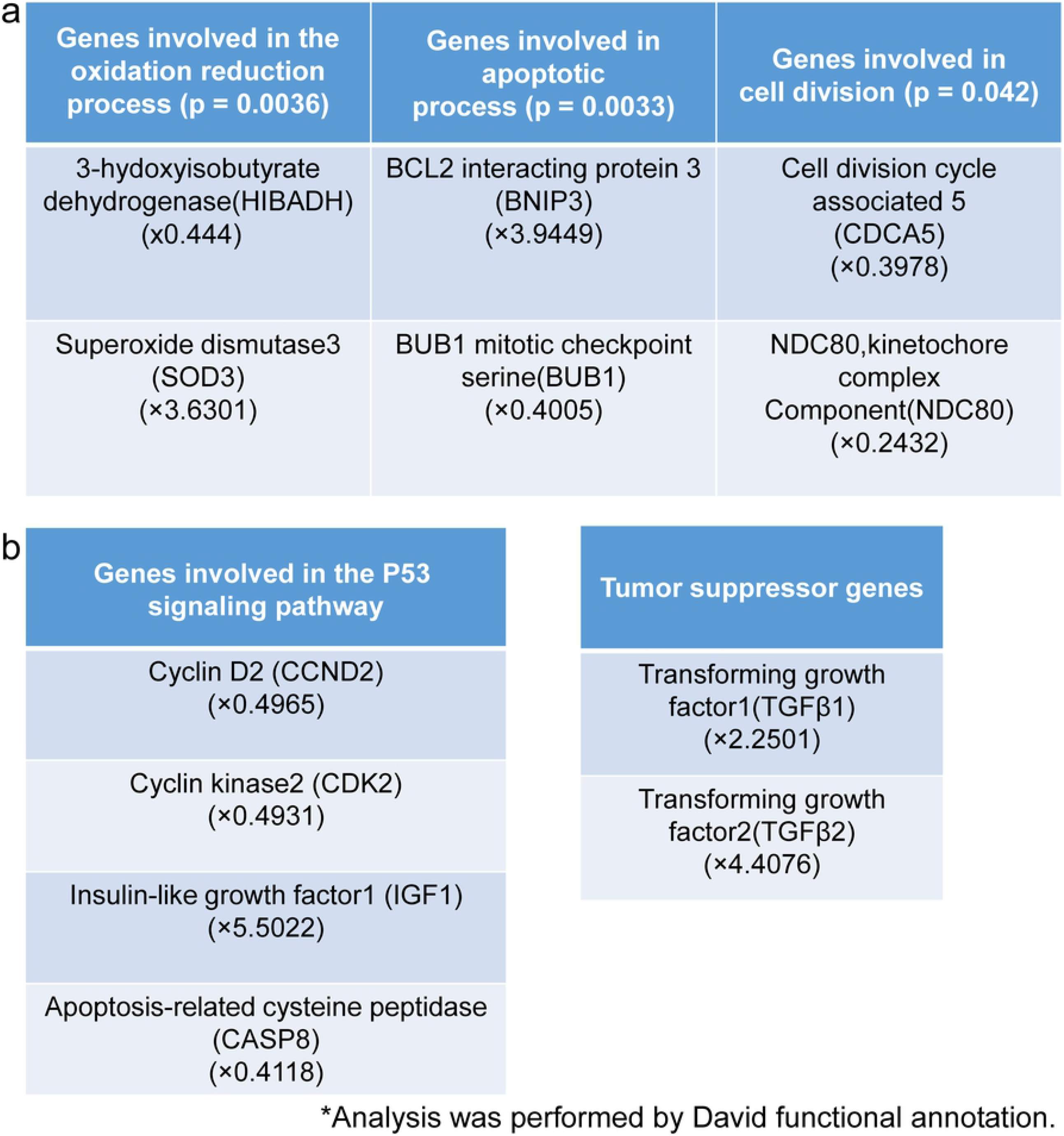
we found that the expression of genes involved in oxidation- reduction and apoptotic processes, and cell division, were significantly changed (p < 0.05).

### Immunohistochemical staining of liver metastases

The median numbers of Ki67-positive cells in each group were compared: HCT116 (control group: 74.3; administration group: 24.3; p = 0.0159); HT29 (control group: 92.3; administration group: 39.3; p = 0.0159); and DLD-1 (control group: 99.7; administration group: 11.0; p = 0.0119). The p-value for the liver metastasis in mice (n = 15) was p = 0.000013. Liver metastasis-positive cells significantly decreased in the antibody-treated groups with each of the cell lines in these animal models.

## Discussion

At present, antibody-based drugs, such as the ones targeting VEGF, are used clinically as molecularly-targeted therapeutic agents for gastrointestinal cancer, and improved prognosis has been observed in unresectable advanced colorectal cancer [17]. VEGF is thought to act on the interstitium around cancer cells and facilitate metastasis. According to several reports, VEGF and hematogenous metastasis are closely related in lung, breast, and renal cancers [18–20]. Thus, drugs affecting VEGF have anticancer effects. For example, bevacizumab has a VEGF-neutralizing effect and normalizes the vasculature, which improves the delivery and effectiveness of chemotherapeutic agents [12-23]. Ziv-Aflibercept is a receptor-antibody complex consisting of VEGF-binding sections from the extracellular domains of VEGF receptors 1 and 2, fused to human immunoglobulin IgG1Fc [24]. Regorafenib targets VEGF 1-3, TIE2, and other receptor tyrosine kinases [25]. However, malignant tumors cannot be suppressed completely via administration of these molecularly targeted therapeutic agents alone; therefore, there is a demand for better therapeutic drugs.

Hematogenous metastases, which involve multiple angiogenic factors, account for about 80% of remote metastases in colorectal cancer. Thus, controlling hematogenous metastases is an important therapeutic strategy for this disease [26,27]. To date, our laboratory has carried out continuous studies focusing on the angiogenic factor, PROK1, as a target molecule for treating cancer. The anti-PROK1 monoclonal antibody produced by our department is a neutralizing antibody that has been proven to inhibit the angiogenic and subcutaneous tumorigenic potentials when added to colorectal cancer cell lines [12,13,15]. While PROK1 staining is not observed in the normal mucosa of the large intestine, it is present in the cytoplasm of cells in about 40% of colorectal cancers. Positive staining is significantly higher in stages III and IV, which indicates more advanced colorectal cancer than stages I and II. It has also been clarified that PROK1 expression is a prognostic factor in colorectal cancer. Specifically, the 5- year survival rate is lower when PROK1 expression is observed than that when it is not observed in stage III and IV colorectal cancer [28].

Goi et al. demonstrated that an anti-PROK1 antibody inhibited angiogenesis and tumorigenic potential [4]. In the present study, we applied our anti-PROK1 antibody in a mouse model of cancer to investigate how it acted on liver metastases. To maintain its concentration in the blood, the anti-PROK1 antibody was administered intraperitoneally the day before tumor implantation and every 3 days thereafter. In the antibody-treated group, liver metastases were decreased and a significant increase in survival time was observed when compared to the control group.

The microarray revealed that genetic changes were suppressed upon administration of the anti-PROK1 antibody; changes were inhibited in 72.1% of the 24,351 analyzed genes. The altered genes included some involved in the p53 cascade as well as an increase in the tumor suppressor gene, TGF-β. TGF-β contributes to growth inhibition, cell differentiation, and the induction of apoptosis in many cells, including epithelial cells. In addition, TGF-β is involved in the epithelial-mesenchymal transition in cancer cells, and enhances the motility and infiltration of epithelial cells [29].

In this study, we examined the action of TGF-β indirectly by performing immunostaining to detect liver metastases in both groups using an anti- Ki67 antibody. The results showed significantly fewer Ki67-positive cells in the antibody-treated group, indicative of inhibited metastases. Overall, the proliferation of colorectal cancer cells was suppressed in the anti- PROK1 antibody-treated group. We believe that administration of the anti- PROK1 antibody suppressed liver metastases by increasing the expression of TGF-β and enhancing cancer cell suppression. Moving forward, we intend to examine the application of the anti-PROK1 antibody in other gastrointestinal cancers using the peritoneal dissemination model, as well as in gastric cancer and pancreatic cancer cell lines.

## Conclusion

We found that the anti-PROK1 antibody acted on the oxidation-reduction and apoptotic processes, and inhibited cell division of tumors in a liver metastasis mouse model using human colon cancer cell lines. We also believe that the increased expression of TGF-β following anti-PROK1 antibody administration may prolong the survival time of mice by suppressing cell growth and liver metastases.

## Notes

### Competing Interest Statement

The authors have declared that no competing interests exist.

